# Methods of inactivation of SARS-CoV-2 for downstream biological assays

**DOI:** 10.1101/2020.05.21.108035

**Authors:** Edward I. Patterson, Tessa Prince, Enyia R. Anderson, Aitor Casas-Sanchez, Shirley L. Smith, Cintia Cansado-Utrilla, Lance Turtle, Grant L. Hughes

## Abstract

The scientific community has responded to the COVID-19 pandemic by rapidly undertaking research to find effective strategies to reduce the burden of this disease. Encouragingly, researchers from a diverse array of fields are collectively working towards this goal. Research with infectious SARS-CoV-2 is undertaken in high containment laboratories, however, it is often desirable to work with samples at lower containment levels. To facilitate the transfer of infectious samples from high containment laboratories, we have tested methods commonly used to inactivate virus and prepare the sample for additional experiments. Incubation at 80°C, and a range of detergents and UV energies were successful at inactivating a high titre of SARS-CoV-2. These protocols can provide a framework for in house inactivation of SARS-CoV-2 in other laboratories, ensuring the safe use of samples in lower containment levels.

## Introduction

The novel coronavirus, severe acute respiratory syndrome coronavirus 2 (SARS-CoV-2) emerged in December 2019 in Wuhan, China, and spread to the rest of the world in a few months causing a pandemic (1,2). This virus causes the coronavirus disease, known as COVID-19, in humans and as of May 17, 2020, has infected almost 5,000,000 people and caused over 300,000 deaths (3). Research on SARS-CoV-2 has increased exponentially since the beginning of the pandemic and will likely continue growing until an effective vaccine is developed. In the UK, and many other countries, SARS-CoV-2 is classified as a hazard group 3 pathogen. For handling clinical samples and performing experiments involving SARS-CoV-2 and other viruses in general, inactivation methods are needed in order to work under safe conditions. Additionally, the inactivation of the virus allows the transfer of the material from a containment level (CL) 3 to a CL2 laboratory, facilitating the performance of experiments and increasing the number of laboratories and researchers that can perform those experiments. Several methods of inactivation are available, but since this is a novel virus, the effectiveness of many of these methods on SARS-CoV-2 has not been tested yet. Some inactivation approaches have been tested on SARS-CoV, a coronavirus which spread between November 2002 and September 2003 and whose genome presents a 80% shared identity with the new SARS-CoV-2 (4). It is expected that the outcome of both physical and chemical inactivation methods used against SARS-CoV-2 will be similar to SARS-CoV, but methods need to be validated prior to use of the new virus isolate.

Several methods for virus inactivation are available and the choice of which approach is used is often related to their compatibility with downstream applications. Heat inactivation has been used for several viruses (5,6) and is a common method employed for antigen preservation of viral and bacterial pathogens. To preserve proteins in the sample that are related to host immune response, detergents and UV can be used to inactivate viruses (7,8). Detergents are common additives in reagents used for virus inactivation, as well as RNA extraction from a range of sample types. UV irradiation, which inactivates viruses by modifying their nucleic acid structure, has been used successfully to inactivate many viruses, and in particular SARS-CoV (9). Inactivation of SARS-CoV-2 through the use of UV would allow the safe use of the virus within a CL2 laboratory and prevent the possibility of lab-acquired infections. Here we aim to assess and describe physical and chemical inactivation protocols of SARS-CoV-2.

## Materials and Methods

### Cell culture and viruses

Vero E6 cells (C1008; African green monkey kidney cells) were obtained from Public Health England and maintained in Dulbecco’s minimal essential medium (DMEM) containing 10% fetal bovine serum (FBS) and 0.05 mg/mL gentamycin at 37°C with 5% CO2. SARS-CoV-2 isolate REMRQ0001/Human/2020/Liverpool was cultured from a clinical sample and passaged four times in Vero E6 cells. The fourth passage of virus was cultured in Vero E6 cells with DMEM containing 4% FBS and 0.05 mg/mL gentamycin at 37°C with 5% CO2 and was harvested 48 hours post inoculation. Virus stocks were stored at −80°C.

### Virus inactivation

All inactivation conditions were performed with 1.145 x 10^7^ plaque forming units (PFU) of virus. Heat inactivation was performed by incubating 300 μl of SARS-CoV-2 stock at 80°C for 1 hour. Inactivation with detergents (0.5% sodium dodecyl sulfate (SDS), 0.5% Triton X-100, 0.5% Tween 20, 0.5% NP-40) were incubated with virus for 30 minutes at room temperature. UV inactivation was performed using a CL1000 UVP Crosslinker (UVP, Upland, CA, USA). The CL-1000 UVP Crosslinker consisted of 5 x 254 nm shortwave tubes, which is the recommended wavelength for neutralising viruses, in particular SARS-CoV (9). Virus stock was added (250 μl) into wells of a 24-well plate and placed without its lid on top of an ice block inside the crosslinker. Plates were placed exactly 6 cm from the UV bulbs. Inactivation was performed at a range of UV energy exposures; 0.01 J/cm^2^ – 0.8 J/cm^2^. All inactivation procedures were performed in triplicate, with the exception of NP-40 which was performed in duplicate.

### Virus viability and quantification

Heat treated samples were evaluated for viable virus in a TCID_50_ assay with Vero E6 cells, using the entire volume of the sample. Control virus stocks containing 10^0^, 10^1^ and 10^2^ PFU incubated at room temperature for 1 hour were used to determine the limit of detection of the assay. TCID_50_ plates were passaged onto fresh cells for 4 days at least twice to ensure no replicative virus remained. Cells were monitored daily for cytopathic effect (CPE).

Inactivation treatments using SDS and Triton X-100 were added to 15 mL of DMEM in a centrifugal concentrator (Amicon Ultra-15 100kDa MWCO) and centrifuged until £300 μl of sample remained. Conditions with Tween 20 and NP-40 were diluted in 50 mL of PBS and concentrated until £300 μl of sample remained. Sample was extracted and virus culture medium was added to a final volume of 300 μl. Viable virus was evaluated in a TCID_50_ assay as outlined above. Control virus stocks containing 10^0^, 10^1^ and 10^2^ PFU were diluted in PBS and followed the above protocol with centrifugal filters to determine the limit of detection of the assay.

Plaque assays and TCID_50_ assays were performed on untreated virus stocks and on UV inactivated stocks in parallel. TCID_50_ plates were passaged onto fresh cells for 3 days at least twice to ensure no replicative virus remained. TCID_50_ results were calculated using the Spearman and Karber method as previously described (10).

## Results

We first determined the limits of sensitivity for our detection method by quantifying SARS-CoV-2 at 10^0^, 10^1^, or 10^2^ PFU using TCID_50_ assays. This was done by quantifying viral titres directly, or by passing the sample through a centrifugal column, which is used to remove the inactivation agent before assaying on cells. Virus prepared without the centrifugal column was detected down to dilutions of 10^0^ PFU of SARS-CoV-2 (Figure 1A). The limit of detection with the centrifugal columns was determined to be 10^1^ PFU of SARS-CoV-2 (Figure 1A).

**Figure 1.**
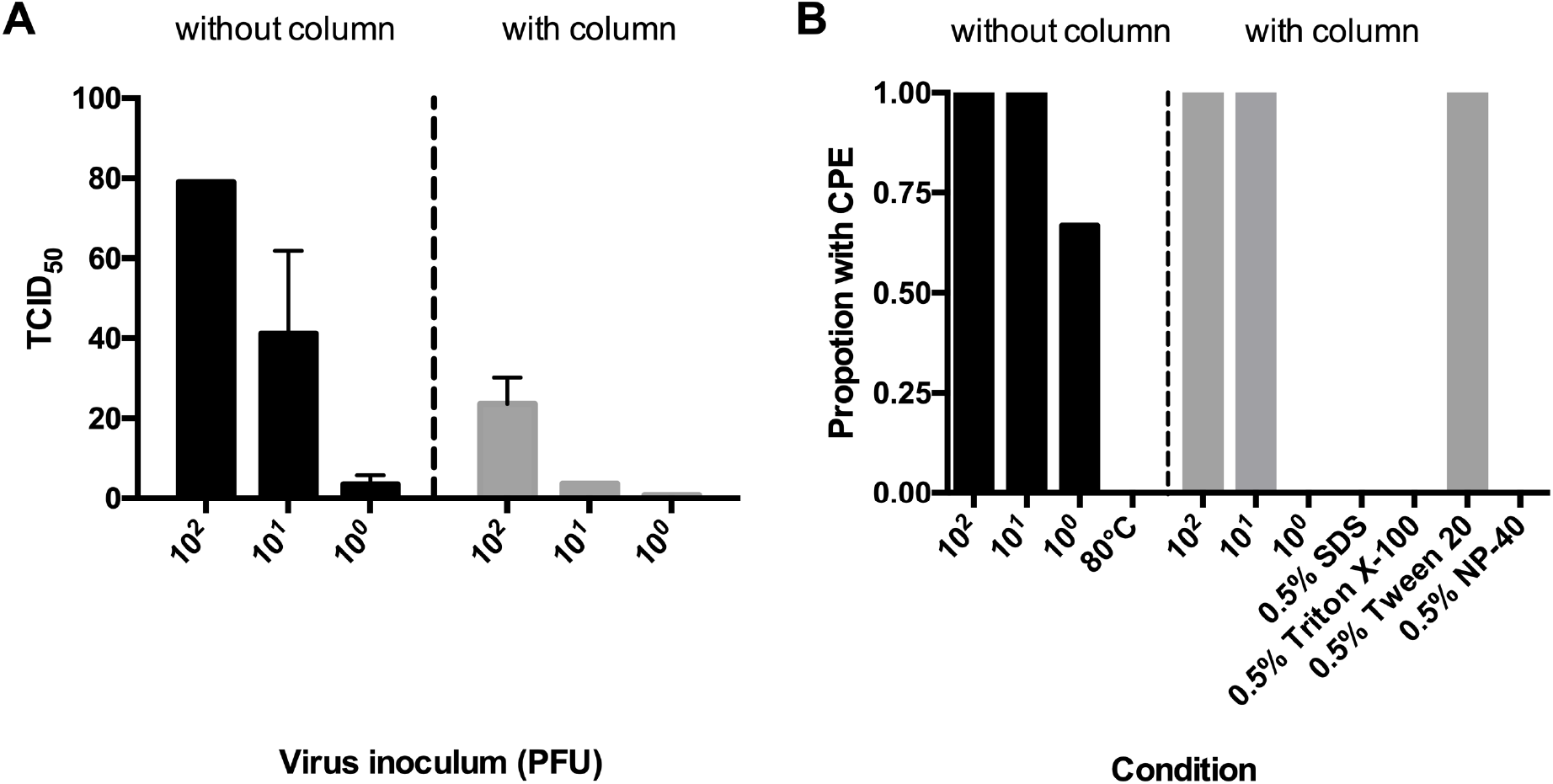
Detection of infectious SARS-CoV-2. A) Assay to quantify the limits of detection. Known titres of virus were prepared with (right) or without (left) concentrating the sample in a centrifugal column. Quantification of controls was performed using TCID_50_ (lower limit of detection is 3.16 TCID_50_/mL). N=3; mean ± SEM. B) The proportion of inactivation assays with cytopathic effect (CPE). Samples were either diluted for assays (right) or inactivation removed using centrifugal columns (right). Control samples with 10^0^, 10^1^, and 10^2^ PFU of SARS-CoV-2 were used as positive controls and to determine the limit of detection for each method.

We then quantified virus after inactivation. SARS-CoV-2 treated at 80°C for one hour was successfully inactivated. We passaged samples a second time to confirmed complete viral inactivation. Using detergents, complete inactivation was seen in all replicates of SDS, Triton

X-100 and NP-40, which we again confirmed by passaging the supernatant to a fresh monolayer of cells to check for residual virus (Figure 1B). However, we found virus samples treated with Tween 20 all remained infectious. At the 10^0^ PFU dilution, CPE was observed in 2 of 3 replicates (Figure 1B), however virus was not detected in this dilution when using the centrifugal columns for clean-up. CPE was seen in all other control wells for either treatment.

In order to determine if UV exposure at 254 nm would inactivate SARS-CoV-2, virus stocks were placed in wells of a 24-well plate placed on ice and exposed to varying amounts of UV energy (J/cm^2^). Exposure of SARS-CoV-2 to UV light at 0.01 J/cm^2^ resulted in partial inactivation and this increased with greater UV energy exposure, resulting in complete inactivation at UV energy exposures of more than 0.04 J/cm^2^ (Figure 2). A similar inactivation curve was seen by both TCID_50_ and plaque assay. No CPE was evident in further passages in samples where inactivation was observed.

**Figure 2.**
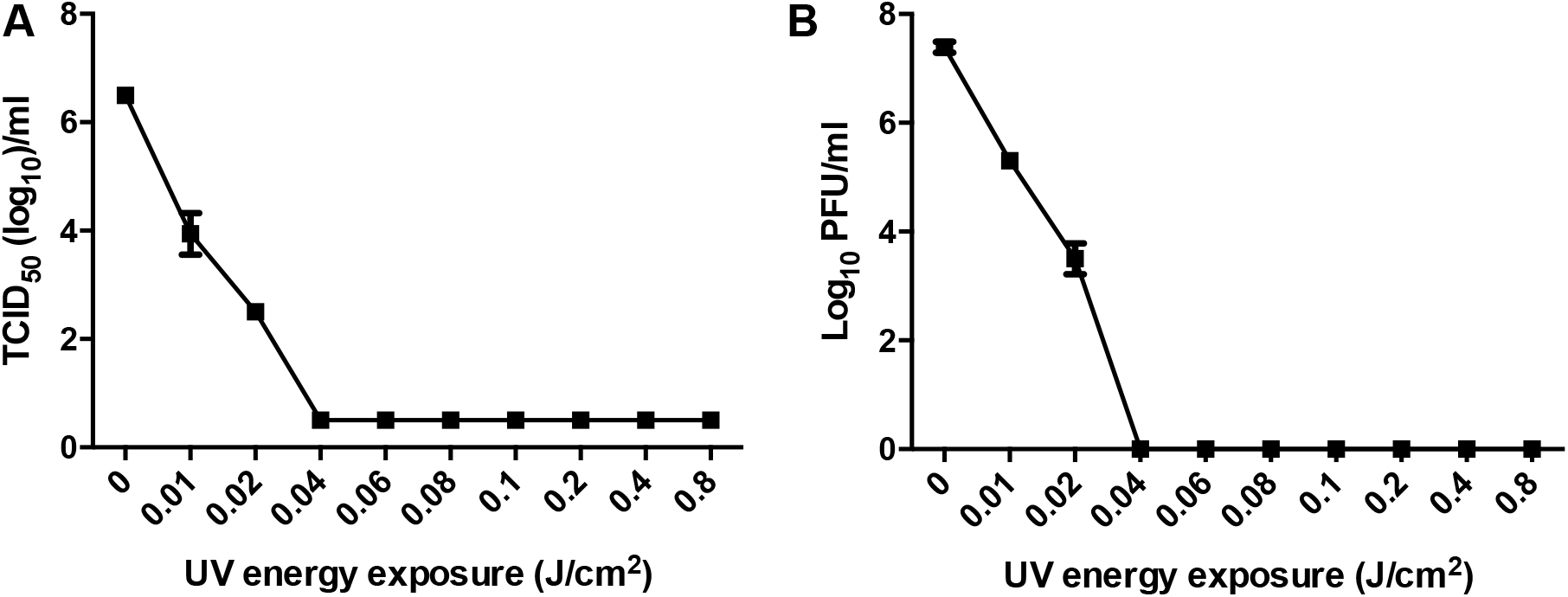
Quantification of SARS-CoV-2 following exposure to different energies of UV. A) The concentration of viable SARS-CoV-2 following exposure to UV measured by TCID_50_ assay (lower limit of detection is 3.16 TCID_50_/mL). B) The concentration of viable SARS-CoV-2 following exposure to UV measured by plaque assay. Both assays confirmed the complete inactivation of the sample occurring at 0.04 J/cm^2^.

## Discussion

Virus inactivation can be achieved by several methods. However, specific methods must be chosen to comply with requirements for subsequent downstream experiments. We used a range of techniques that are often used for preserving antigens or immunological proteins to evaluate their ability to inactivate SARS-CoV-2, including a range of common detergents and determining the threshold of inactivation by UV exposure. The assay to assess infectious particles was also shown to be sensitive, in some cases down to a single infectious virus particle, and down 10 infectious particles/mL where a centrifugal column is used to concentrate the sample.

Physical inactivation can be performed using heat or exposure to UV. Heat inactivates the virus by denaturing the structure of the proteins, affecting the attachment and replication of the virus in the host cell (11). In this study, SARS-CoV-2 was successfully inactivated with a temperature of 80°C. Lower temperatures used to inactivate SARS-CoV showed that 56°C is only effective in the absence of fetal calf serum and temperatures up to 75°C are needed for successful inactivation of infected clinical samples (9,12). However, heat inactivation is not recommended in a clinical setting for immunological assays since it can interfere with the analysis of antibodies against SARS-CoV-2 (13) and diagnosis of patient samples using RT-PCR, which could potentially lead to a false negative diagnosis (14,15).

UV light causes genetic damage by inducing pyrimidine dimers or by producing reactive oxygen species (16). While other investigators have investigated UV inactivation of viruses by looking at length of exposure, here we have inactivated virus based on the energy exposure. As UV lamps age, their irradiance output begins to decline. The crosslinker in this study has an inbuilt sensor allowing the unit to determine the exact amount of UV energy delivered. Therefore, to maintain consistency in experiments over time, it is recommended to inactivate virus based on the UV energy exposure rather than time of exposure. UVC exposure at 3 cm for 15 minutes has been shown to inactivate SARS-CoV, whereas UVA light was not effective (9,17). Here, we have demonstrated a method by which SARS-CoV-2 can be rendered non-infectious through application of UV energy >0.04 J/cm^2^.

Chemical inactivation can be performed using detergents, and we successfully demonstrated this with three different compounds: 0.5% SDS, 0.5% Triton X-100 and 0.5% NP-40. Conversely, Tween 20 was unable to inactivate SARS-CoV-2 under the same conditions. Detergents disrupt the lipid coat of enveloped viruses and are often present in lysis buffers of commercial nucleic acid extraction kits. These detergents typically do not affect proteins so they can be used in downstream procedures preserving their native structure. Our findings are consistent with previous studies showed that 0.1% SDS with 0.1% NP-40 (9) and 0.3% tri(n-butyl)phosphate (TNBP) with 1.0% Triton X-100 (8) could inactivate SARS-CoV. Recent studies on SARS-CoV-2, showed that several lysis buffers from extraction kits like ATL (1-10% SDS) and VXL (30-50% guanidine hydrochloride and 1-10% Triton X-100) from Qiagen (14) and others containing guanidine hydrochloride (18) and guanidinium (19) inactivated the virus. Several RNA extraction kits contain a lysis buffer effective at inactivating SARS-CoV-2 (20). This is convenient for downstream experiments like qRT-PCR, used for diagnosis. However, not all the laboratories may have access to these kits. The use of centrifugal columns to remove cytotoxic compounds has also been successfully employed in this study, correlating to previous results (5,21); however this raises the threshold of detection by approximately 10 fold.

With the increasing interest in COVID-19, many researchers are now applying their knowledge and expertise to different topics to address this global problem. However, not all researchers have access to containment facilities and essential equipment is not often available at biosafety levels required to work safely with SARS-CoV-2. The inactivation methods described here will contribute to help to diverse research groups to perform their downstream work on SARS-CoV-2.

## Acknowledgements

EIP and ACS were supported by the Liverpool School of Tropical Medicine Director’s Catalyst Fund award. GLH was supported by the BBSRC (BB/T001240/1), the Royal Society Wolfson Fellowship (RSWF\R1\180013), NIH grants (R21AI124452 and R21AI129507), and the NIHR (NIHR2000907). GLH, TP, and LT are both affiliated to the National Institute for Health Research Health Protection Research Unit (NIHR HPRU) in Emerging and Zoonotic Infections at University of Liverpool in partnership with Public Health England (PHE), in collaboration with Liverpool School of Tropical Medicine and the University of Oxford. GLH is based at LSTM and TP and LT at the University of Liverpool. The views expressed are those of the author(s) and not necessarily those of the NHS, the NIHR, the Department of Health or Public Health England. LT is supported by a Wellcome clinical career development fellowship, grant number 205228/Z/16/Z and SLS is supported by the DogsTrust. We thank Tom Curry and David Matthews for their gift of the SARS-CoV-2 isolate.

